# Molecular and spatial characterization of baicalin from *Scutellaria baicalensis* hairy root culture

**DOI:** 10.64898/2026.05.20.726740

**Authors:** Fedorova Anastasia Mikhailovna, Milentyeva Irina Sergeevna, Asyakina Lyudmila Konstantinovna, Prosekov Alexander Yuryevich

## Abstract

This study presents the structural verification of baicalin isolated from a hydroethanolic extract of an *in vitro Scutellaria baicalensis* root culture using X-ray diffraction analysis and a set of NMR spectroscopy techniques. The crystalline molecular structure of the sample was found to correspond to baicalin. The ^1^H, ^13^C{^1^H}, 2D ^1^H^1^H-COSY, ^1^H^13^C-HSQC, ^1^H^13^C-HMBC spectra confirmed that the chemical shifts, signal multiplicities, integral intensities, and spin–spin coupling constants were fully consistent with the structure of the target compound. Minor impurity signals were detected in the aliphatic region of the spectra, with a total content not exceeding 5 mol%. These results confirm the high purity and structural individuality of baicalin, a biologically active flavonoid glycoside of considerable interest.

## Introduction

Research aimed at isolating and studying individual biologically active compounds (BAC) from medicinal plant materials and plant cultures obtained by biotechnological methods is currently of particular relevance [1]. This growing interest is due to the fact that plant-derived phenolic metabolites exhibit a wide range of biological activities and are regarded as potential components of medicinal products and functional foods [2]. Among these compounds, flavonoids occupy a special place because of their wide occurrence and important role in plant metabolism.

Baikal skullcap *(Scutellaria baicalensis* Georgi) is a valuable medicinal plant traditionally used in medicine and a source of various biologically active compounds. Phenolic constituents are especially important among them, including baicalin, a flavonoid glycoside with pronounced biological activity [3]. Antioxidant [4], anti-inflammatory [5], antibacterial, and neuroprotective properties [6] have been reported for baicalin, which accounts for the continuing interest in its isolation, purification, and structural identification.

*Scutellaria baicalensis* hairy root cultures are regarded as one of the most promising systems for baicalin production because they show stable growth and are capable of accumulating the target secondary metabolites [7]. The use of such cultures makes it possible to obtain standardized natural compounds independently of the seasonal and environmental factors typically associated with plant raw materials. In this context, a comprehensive study of the molecular and spatial structure of baicalin isolated from *S. baicalensis* hairy root biomass is of particular relevance.

Biologically active compounds in medicinal plant materials are most commonly identified by liquid chromatography combined with various detection methods, including ultraviolet and infrared spectroscopy, nuclear magnetic resonance (NMR) spectroscopy, and mass spectrometry (MS) [8]. However, despite the widespread use of these approaches, relatively few studies have focused on the comprehensive structural verification of natural compounds, and in some cases the reported results remain ambiguous or contradictory. Therefore, methods that not only confirm the composition of a substance but also clarify the features of its molecular and spatial structure are of particular importance.

For confirming the structure of natural compounds, the combined use of NMR spectroscopy and X-ray diffraction analysis is considered the most informative approach, as these methods make it possible not only to establish the molecular structure but also to characterize in detail the spatial organization of the molecule, its stereochemical features, and its crystal packing [9]. NMR spectroscopy provides information on the nature of chemical bonds, the relative arrangement of atomic fragments, and the types of functional groups present in solution, whereas X-ray diffraction analysis offers a direct description of the three-dimensional structure of a compound in the crystalline state [10]. The combined use of these methods increases the reliability of structural identification, particularly in studies of natural compounds with closely related structures and possible accompanying impurities.

The advantages of NMR over MS, including MS used in combination with gas or liquid chromatography, include high reproducibility, minimal sample preparation, the non-destructive nature of the analysis, and the ability to obtain quantitative information without the mandatory use of reference standards [11]. In addition, NMR is particularly informative for the analysis of highly polar compounds, which may be difficult to determine by MS. At the same time, MS offers higher sensitivity and selectivity, making it effective for the analysis of complex multicomponent mixtures [12, 13]. Thus, NMR and MS are complementary methods, and their combined use substantially improves the reliability of structural identification of natural compounds.

The aim of this study was to investigate the molecular and spatial structure of baicalin isolated from *Scutellaria baicalensis* hairy root culture using NMR spectroscopy and X-ray diffraction analysis.

## Materials and Methods

The object of the study was baicalin obtained from a hydroethanolic extract of *S. baicalensis* hairy root culture.

Baicalin isolated from the extract of *Scutellaria baicalensis* root culture had been obtained at earlier stages of the study. The methods used for cultivating *Scutellaria baicalensis* root culture and for extracting *S. baicalensis* hairy root biomass are described in the study by A. M. Fedorova and colleagues [14]. Seedlings grown for 14–28 days on a nutrient medium of the following composition were transformed using the soil agrobacterial strain *Agrobacterium rhizogenes* 15834 Swiss (Moscow, Russia): B5 macronutrients, 50.00 mg; B5 micronutrients, 10.00 mg; Fe-EDTA, 5.00 mL; thiamine, 10.00 mg; pyridoxine, 1.00 mg; nicotinic acid, 1.00 mg; sucrose, 30.00 g; inositol, 100.00 mg; 6-benzylaminopurine, 0.05 mg; indole-3-acetic acid, 1.00 mg; and agar, 20.00 g. For *S. baicalensis* root cultures *in vitro*, the cultivation cycle was 5 weeks. The hydroethanolic extract of *S. baicalensis* root culture was prepared using 30±0.2 ethanol at a raw material-to-extractant ratio of 1:86, at 70±0.1 °C for 6±0.1 h.

The extraction and purification of baicalin from the *S. baicalensis* root culture extract are described in the study by E. R. Faskhutdinova and colleagues [15]. The purification scheme for baicalin obtained from the *S. baicalensis* root culture extract is shown in Figure 1. The purity of baicalin was at least 95%.

**Figure 1.**
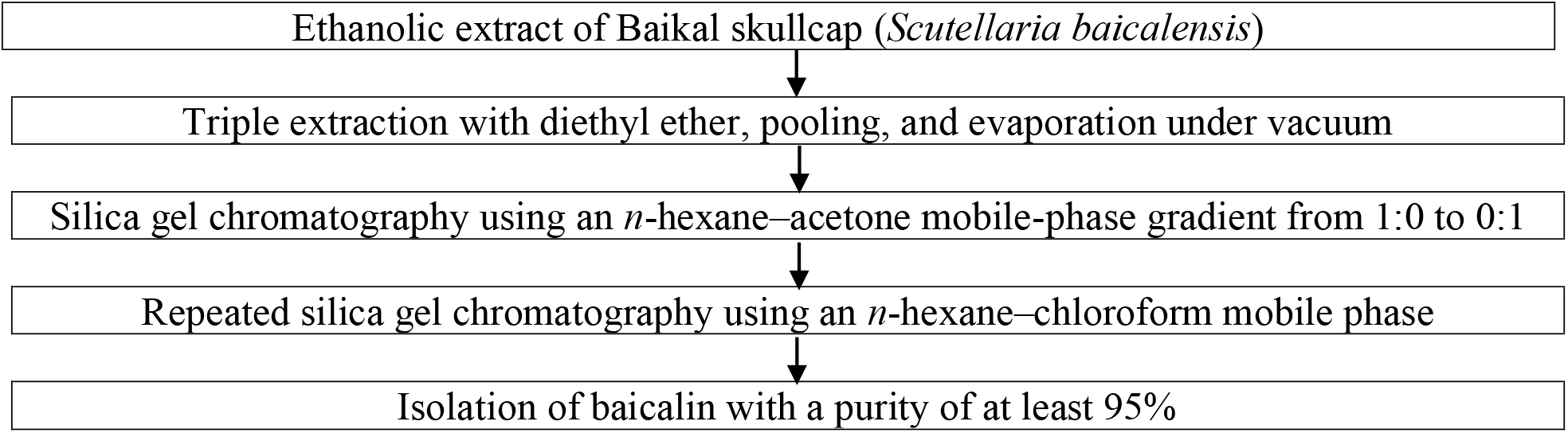
Purification scheme for baicalin obtained from an extract of *Scutellaria baicalensis* root culture Table 1. Operating frequencies used in the study

The crystal structure of baicalin was analyzed using a Bruker D8 Advance X-ray diffractometer with CuKα radiation, equipped with a LynxEye position-sensitive detector. Measurements were performed in reflection geometry with sample rotation. Data were collected using the Bruker DIFFRACplus software package and analyzed with EVA and TOPAS V5.0.

**Table 1.**
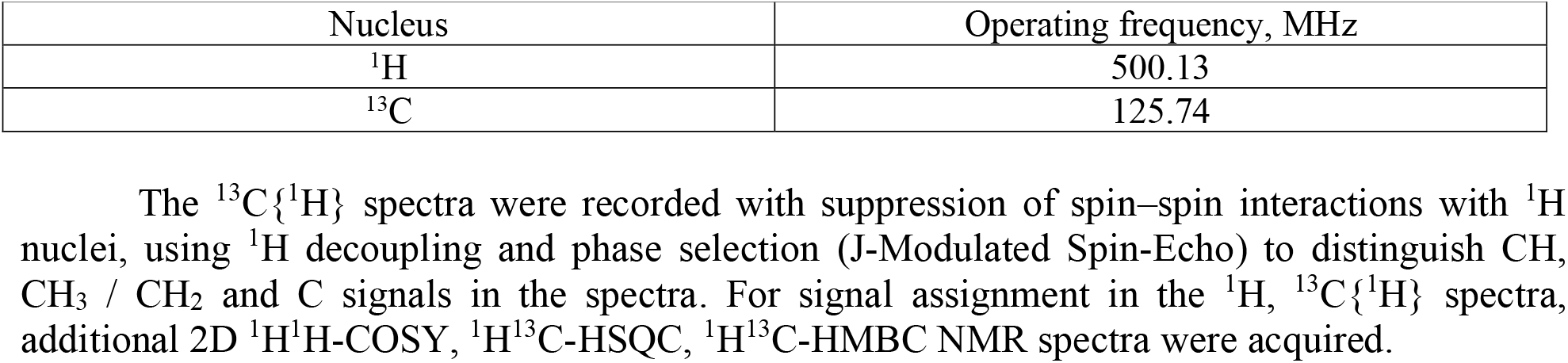
Operating frequencies used in the study.

For NMR spectroscopy, 20 mg portions of the baicalin sample were dissolved in 600 μL of dimethyl sulfoxide (DMSO) at 25 °C. The resulting solutions were placed in a Bruker Avance III HD 500 NMR spectrometer for spectral acquisition. The operating frequencies of the NMR spectrometer are shown in Table 1.

The ^13^C{^1^H} spectra were recorded with suppression of spin–spin interactions with ^1^H nuclei, using ^1^H decoupling and phase selection (J-Modulated Spin-Echo) to distinguish CH, CH_3_ / CH_2_ and C signals in the spectra. For signal assignment in the ^1^H, ^13^C{^1^H} spectra, additional 2D ^1^H^1^H-COSY, ^1^H^13^C-HSQC, ^1^H^13^C-HMBC NMR spectra were acquired.

## Results and Discussion

The analysis showed that the baicalin sample was crystalline and single-phase, as shown in Figure 2 and Table 2.

**Figure 2.**
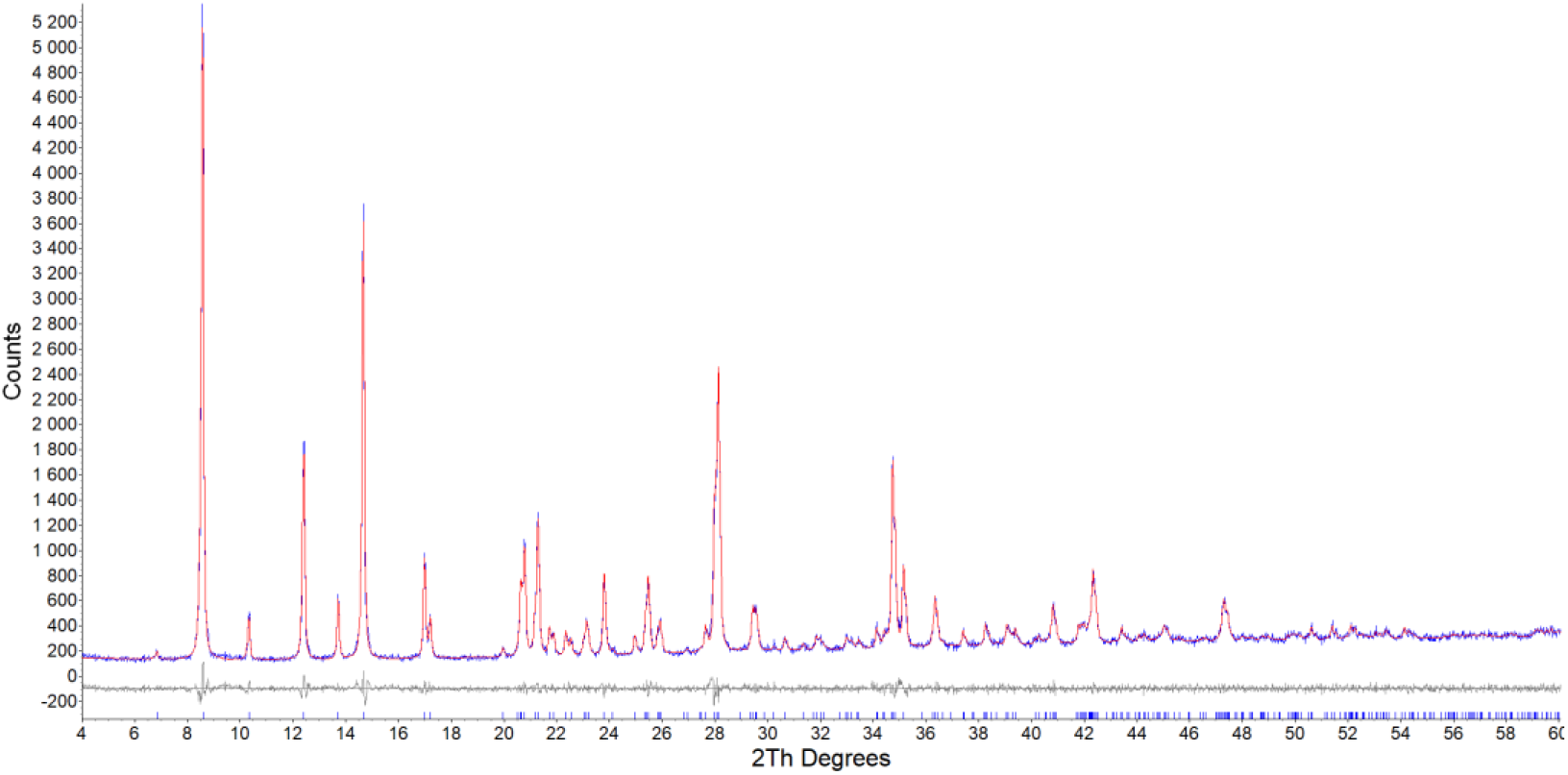
Experimental diffractogram of the baicalin sample (blue curve), theoretical diffractogram calculated for the corresponding unit cell (red line), and the difference curve between them (gray line). Vertical ticks indicate the calculated peak positions.

**Table 2.**
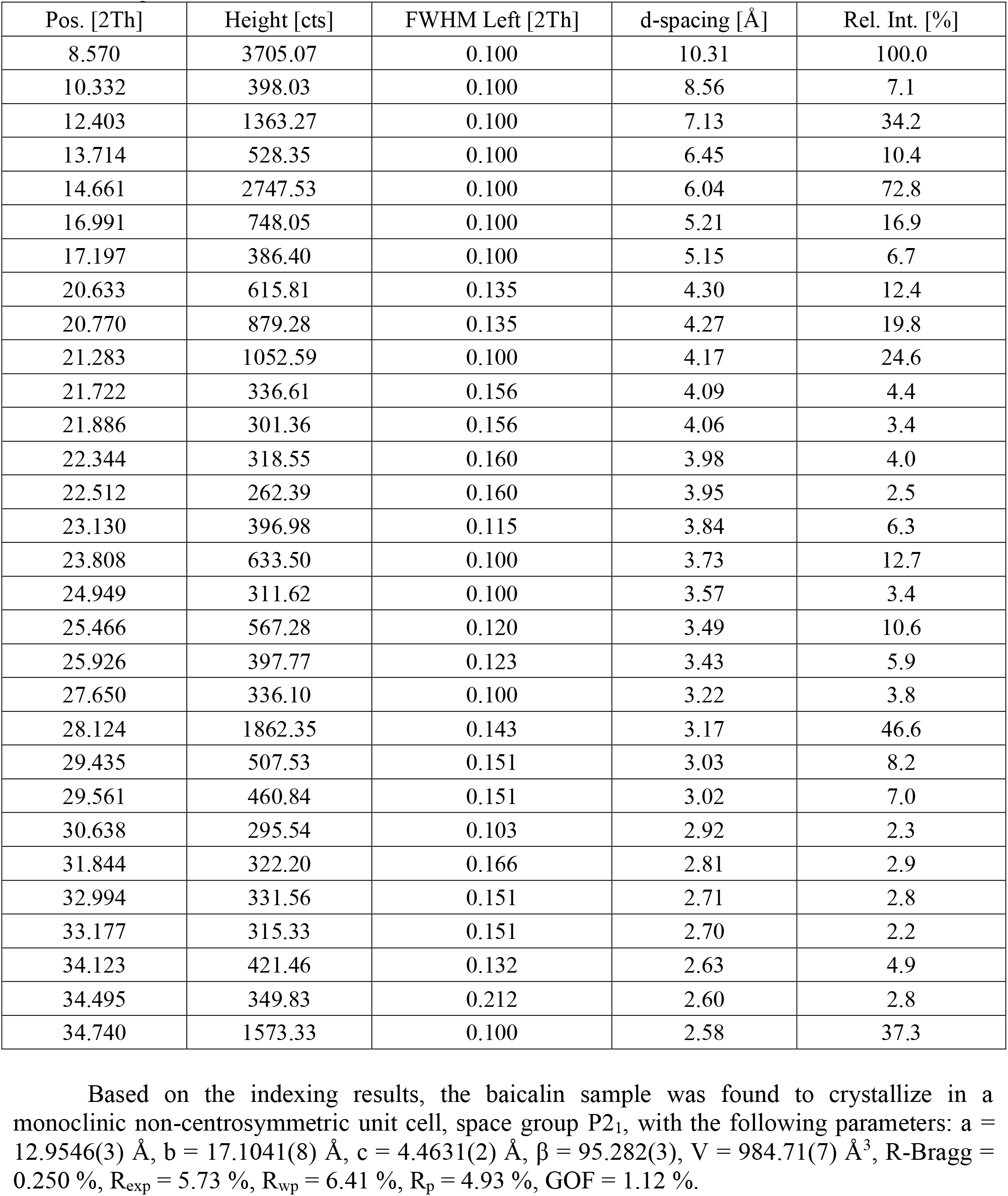
Intensities and positions of selected peaks with a relative intensity above 2% for the baicalin sample.

Based on the indexing results, the baicalin sample was found to crystallize in a monoclinic non-centrosymmetric unit cell, space group P2_1_, with the following parameters: a = 12.9546(3) Å, b = 17.1041(8) Å, c = 4.4631(2) Å, β = 95.282(3), V = 984.71(7) Å^3^, R-Bragg =0.250 %, R_exp_ = 5.73 %, R_wp_ = 6.41 %, R_p_ = 4.93 %, GOF = 1.12 %.

The diffraction analysis showed that the baicalin sample under study corresponded to the declared compound, baicalin.

Additional structural studies of the baicalin sample were then performed using NMR spectroscopy.

Figure 3 shows the NMR spectra of the baicalin sample, which was presumed to have the structure of baicalin: ^1^H, ^13^C{^1^H}, 2D ^1^H^1^H-COSY, ^1^H^13^C-HSQC, ^1^H^13^C-HMBC spectra.

**Figure 3.**
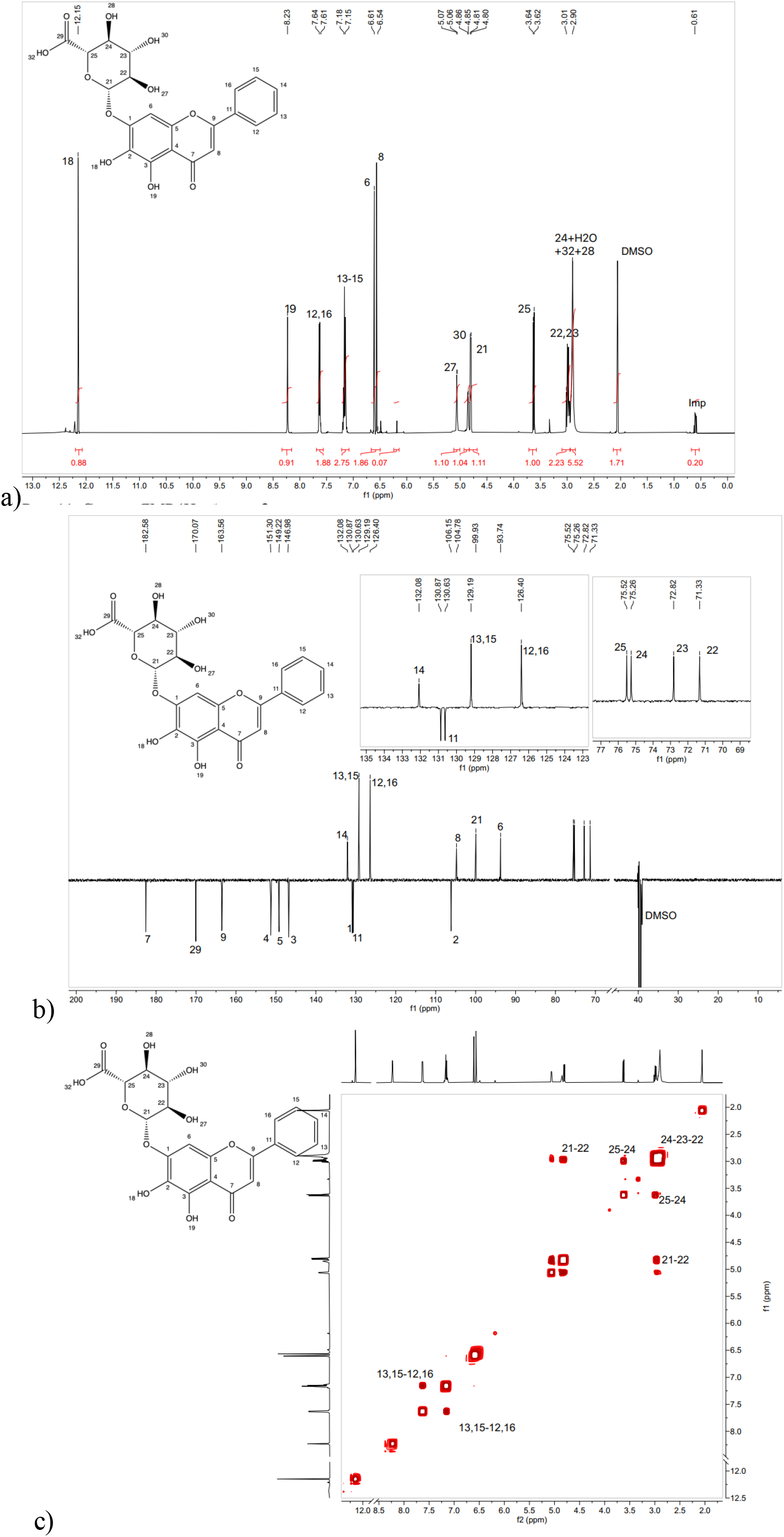
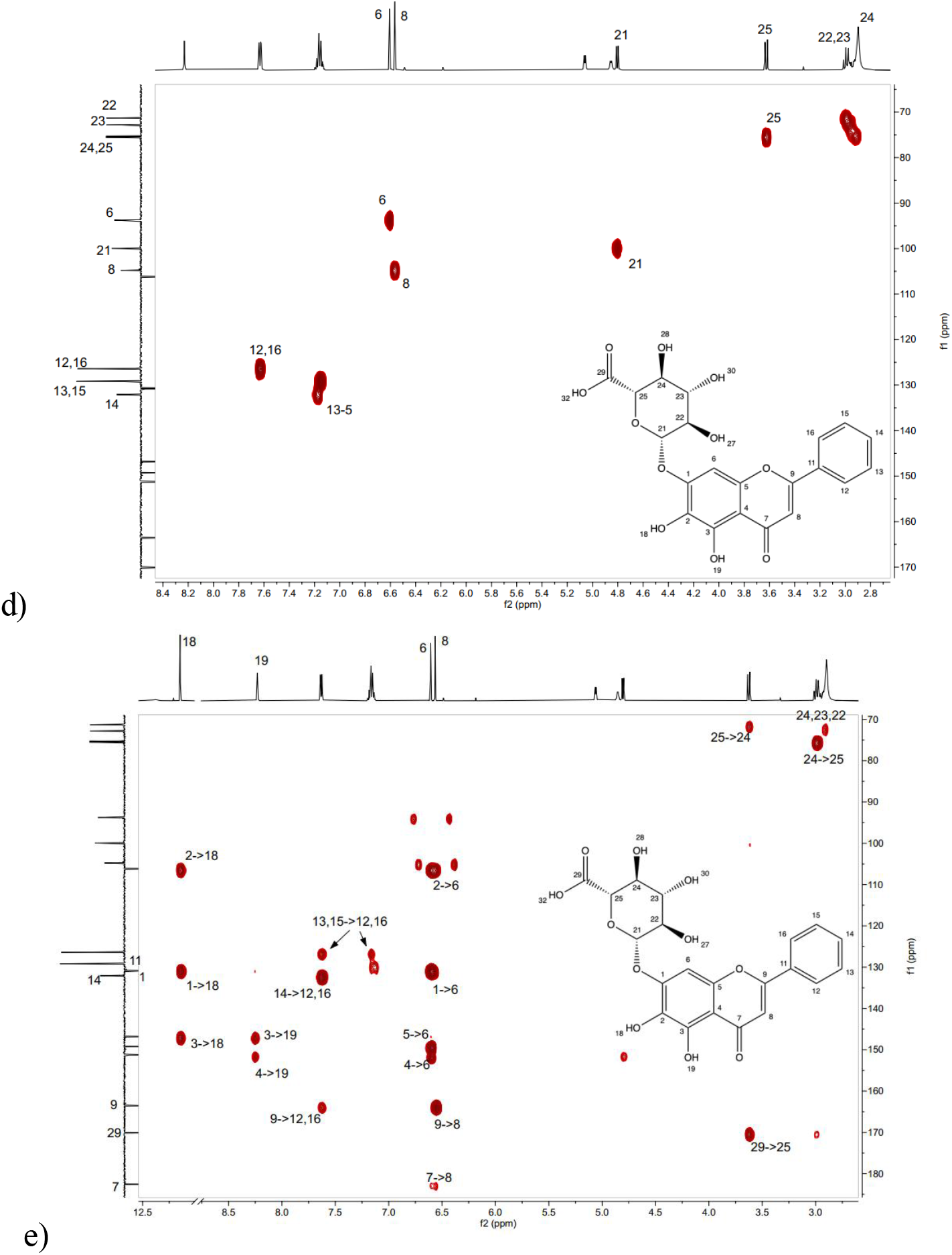
NMR spectra of the baicalin sample: a –^1^H; b –^13^C{^1^H}; c –2D ^1^H^1^H-COSY; d –^1^H^13^C-HSQC; e – ^1^H^13^C-HMBC

The results presented above show that the ^1^H, ^13^C{^1^H}, 2D ^1^H^1^H-COSY, ^1^H^13^C-HSQC, ^1^H^13^C-HMBC NMR spectra of the baicalin sample correspond to the target product, baicalin (Figure 3). Minor impurity signals were present in the aliphatic region of the spectrum, not exceeding 5 mol %. In terms of the number of signals, their positions in the spectra, namely their chemical shifts, integral intensities and ratios, signal multiplicities, and spin–spin coupling constants, the acquired spectra were consistent with the target product (Figure 4).

**Figure 4.**
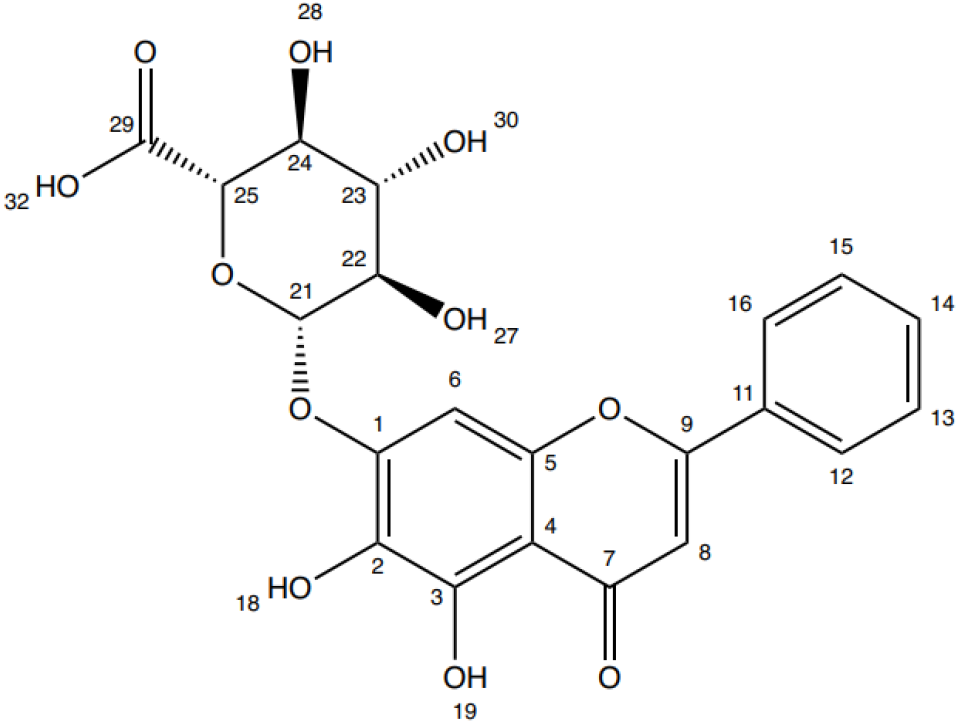
Structural formula of the baicalin sample identified as baicalin.

Signal assignments are presented in Table 3 for the ^1^H and ^13^C{^1^H} spectra.

**Table 3.**
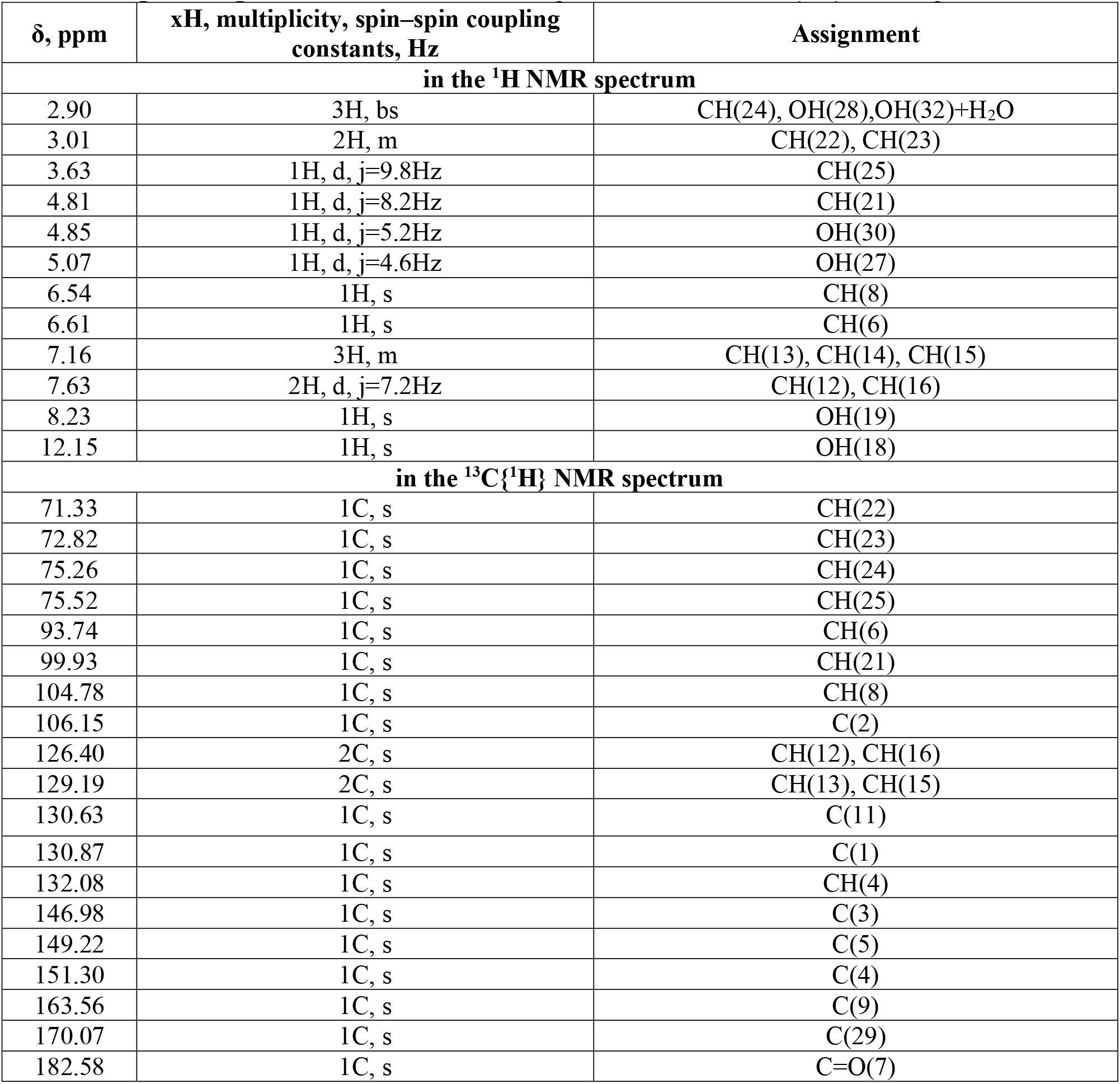
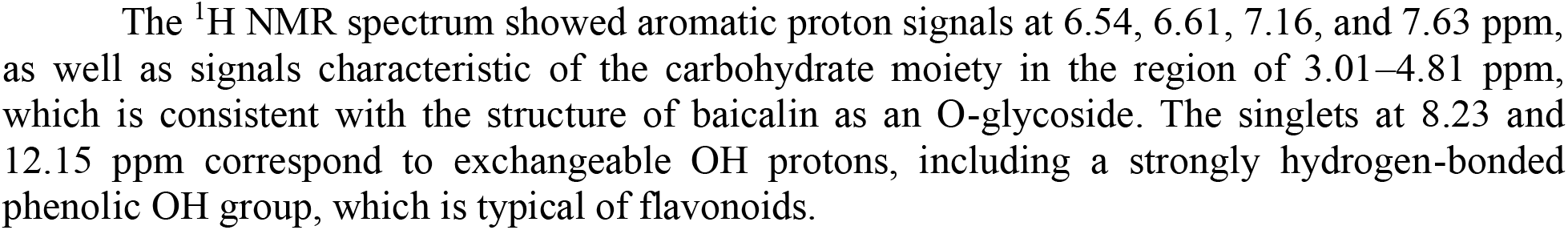
Signal assignments for the baicalin sample in the ^1^H and ^13^C{^1^H} NMR spectra.

The ^1^H NMR spectrum showed aromatic proton signals at 6.54, 6.61, 7.16, and 7.63 ppm, as well as signals characteristic of the carbohydrate moiety in the region of 3.01–4.81 ppm, which is consistent with the structure of baicalin as an O-glycoside. The singlets at 8.23 and 12.15 ppm correspond to exchangeable OH protons, including a strongly hydrogen-bonded phenolic OH group, which is typical of flavonoids.

Minor impurity signals were present in the aliphatic region of the spectrum in the baicalin sample, with a content of up to 5 mol %.

The NMR spectroscopy results confirmed that the study object, baicalin isolated from the hydroethanolic extract of *in vitro S. baicalensis* root culture, corresponded to the declared compound, baicalin.

## Conclusion

The results obtained confirm that *S. baicalensis* root culture is a promising source of baicalin. The use of X-ray diffraction analysis and NMR spectroscopy supports the potential of biotechnologically produced root biomass for the reproducible production and further study of baicalin as a biologically active compound.

## Funding

*This study was carried out within the framework of the state assignment “Development of biologically active supplements composed of metabolites from in vitro plant objects to protect the population from premature aging” (project FZSR-2024-0008)*

*The study was performed using the equipment of the Core Facilities “Instrumental Methods of Analysis in Applied Biotechnology” at Kemerovo State University*.

